# jaxon: a differentiable, GPU-native simulator for peripheral-nerve fiber models

**DOI:** 10.64898/2026.07.30.741846

**Authors:** David Lung, Max Haberbusch

## Abstract

*jaxon* is an open-source, fully differentiable and GPU-native reimplementation of the canonical peripheral-nerve fiber models in JAX/Jaxley: the myelinated McIntyre–Richardson–Grill (MRG) and Sweeney axons and the un-myelinated Sundt and Rattay C-fibers. It reproduces NEURON’s `extracellular` mechanism through a custom backward-Euler coupled intracellular/periaxonal double-cable solver, agreeing with PyFibers-wrapped NEURON on 99.6% of 943 activation-threshold configurations within 1% and matching conduction velocity to machine precision. Because the entire forward model is expressed in JAX, it is both vectorized—simulating whole fiber populations in parallel and reaching a geometric-mean ∼820× speedup at *N* = 100,000 fibers on a single GPU—and differentiable, so extracellular-stimulation parameters (per-contact amplitudes, waveform shape, and electrode position) can be optimized directly through the cable equation rather than grid-searched. *jaxon* slots into existing peripheral-nerve modeling pipelines as a gradient-enabled, population-scale replacement for the NEURON forward solver.

## Required metadata

### 1. Motivation and significance

Electrical stimulation of peripheral nerves is an established therapy for a growing range of conditions, and selecting where to place an electrode and how to shape the stimulus increasingly relies on computational models. The standard modeling approach couples a volume-conductor description of the tissue—typically solved by the finite-element method (FEM)—to biophysical, multi-compartment cable models of individual axons [2, 3]. The extracellular potential computed along each fiber drives the membrane model, and threshold and recruitment follow. The reference implementation of the fiber side of this pipeline is NEURON [7], most conveniently accessed through PyFibers [8], which wraps the canonical published axon models.

**Table 1.**
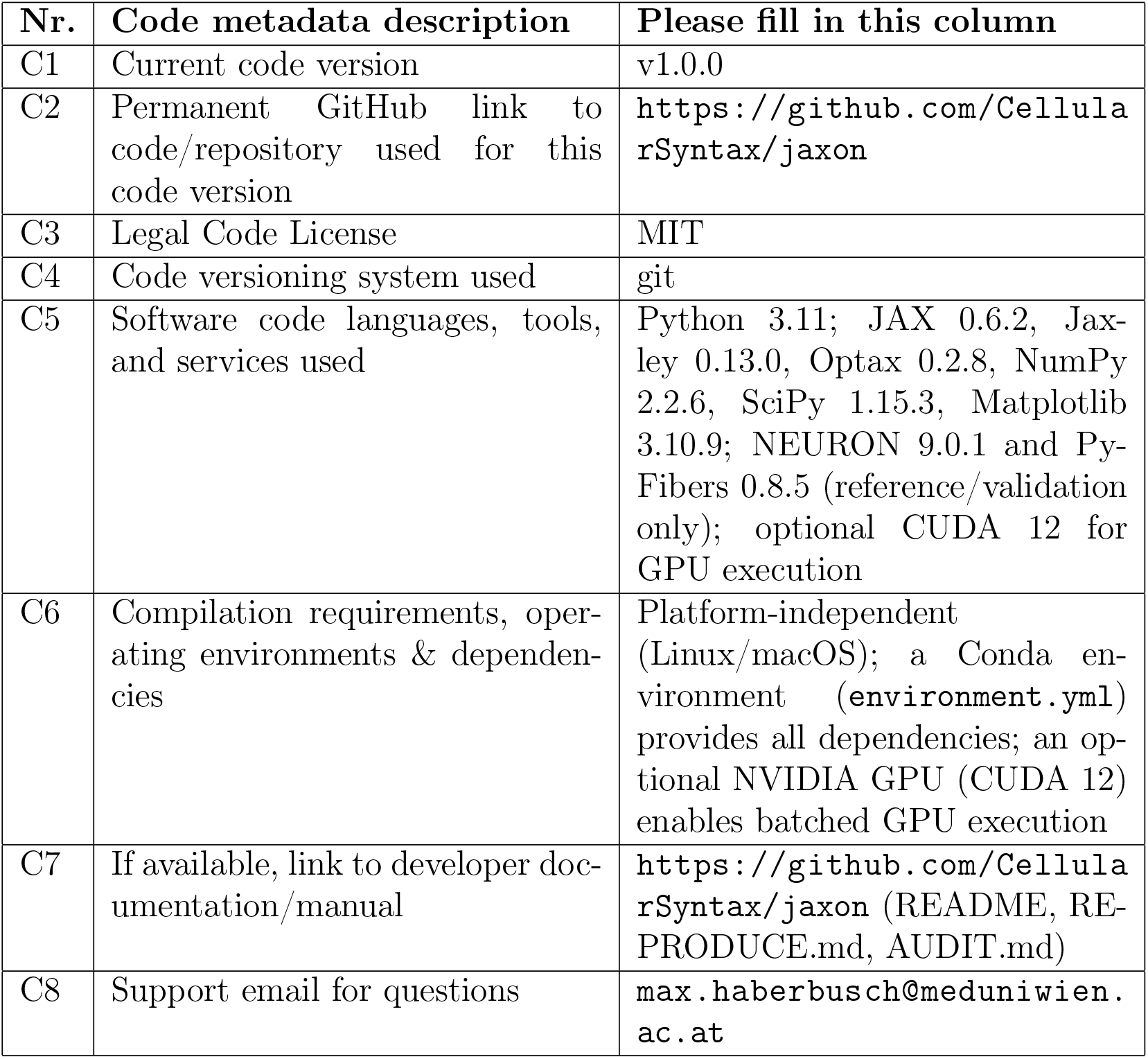
Code metadata.

This reference stack is accurate but has two structural limitations for modern neural-interface design. First, it is *serial* : NEURON simulates one fiber at a time on the CPU, so scoring an optimized stimulus over an entire fiber population—hundreds to tens of thousands of axons per nerve—is prohibitively slow. To stay tractable, models almost universally reduce each fascicle to one or a few representative fibers, and whether that reduction biases the selectivity they predict has been effectively untestable. Second, it is *non-differentiable*: because there is no gradient of the fiber response with respect to the stimulus, stimulation parameters must be tuned by grid search or gradient-free optimization, both of which scale poorly as the number of contacts and waveform degrees of freedom grows. GPU-accelerated surrogates such as AxonML [11] address the speed problem by learning a fast approximation of the threshold, but a surrogate is a fit to the biophysics rather than the biophysics itself.

*jaxon* was built to remove both limitations while keeping the biophysics exact. It reimplements the canonical peripheral-nerve fiber models—the myelinated MRG [3] and Sweeney [4] axons and the unmyelinated Sundt [5] and Rattay [6] C-fibers—in JAX [9] on top of the Jaxley differentiable cable simulator [10], and adds a custom backward-Euler solver for the coupled intracellular/periaxonal double-cable equation that is the principled equivalent of NEURON’s `extracellular` mechanism. Because the whole forward model is JAX, it is simultaneously *vectorized* —running whole populations in parallel with `jax.vmap`—and *differentiable*—exposing exact gradients of the fiber response with respect to the stimulus. *jaxon* is validated against PyFibers-wrapped NEURON and reaches a geometric-mean ∼820× speedup at *N* = 100,000 fibers on one GPU, fast enough to optimize and then re-score whole-nerve populations. A companion paper [1] uses *jaxon* to show that the common single-representative-fiber reduction overestimates deliverable selectivity in a species-dependent way; the present paper describes the software itself.

### 2. Software description

#### 2.1. Software architecture

*jaxon* is organized as a differentiable forward model wrapped by an optimization and analysis layer (Fig. 1). At its center is a vectorized integrator of the coupled (*V*_*i*_, *V*_*px*_) cable equation that maps a set of per-contact stimulus amplitudes (and, optionally, waveforms and electrode positions) onto the membrane response of an arbitrary number of fibers at once. The model is used through three interchangeable entry points that all act on the same underlying functions: a Python package (`jaxon`) for library use, a set of experiment scripts (`experiments_v2`) that reproduce the paper, and SLURM batch drivers for cluster runs.

**Figure 1.**
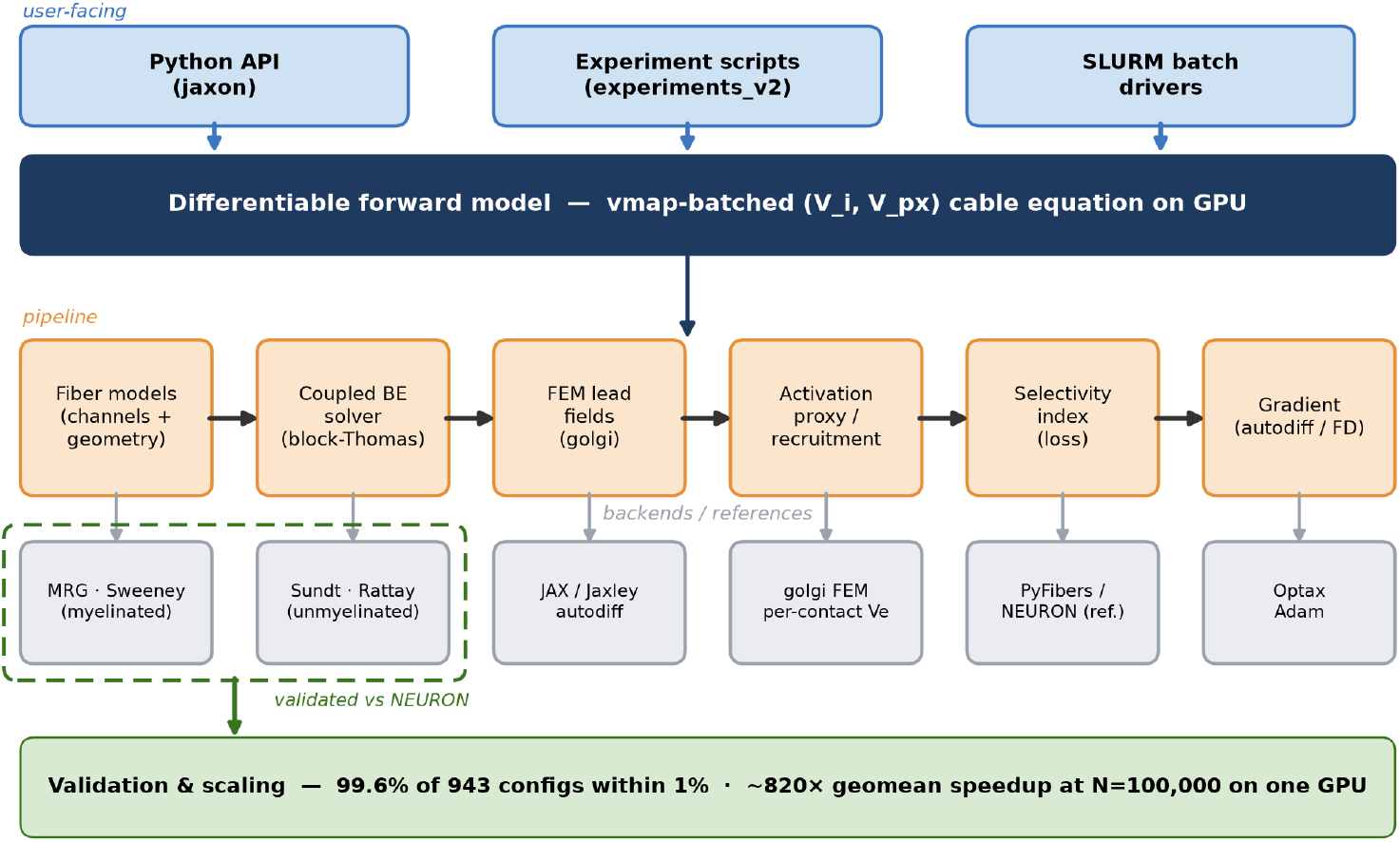
Architecture of *jaxon*. Three user-facing entry points (the `jaxon` Python package, the `experiments_v2` scripts, and SLURM batch drivers) share one differentiable, `vmap`-batched forward model of the coupled (*V*_*i*_, *V*_*px*_) cable equation. The pipeline runs on JAX/Jaxley (autodiff), *golgi* FEM per-contact lead fields, and PyFibers/NEURON as the validation reference; the myelinated (MRG, Sweeney) and unmyelinated (Sundt, Rattay) fiber models, the block-Thomas coupled solver, the activation/selectivity losses, and the gradient form the stages. The whole model is validated against NEURON and scales to *N* = 100,000 fibers on one GPU.

The package is structured by concern:

- `jaxon.channels` — each NMODL ion-channel mechanism hand-translated into a pure-JAX `Channel` class, with rate constants checked against the published `.mod` formulas to machine precision;
- `jaxon.fibers` — morphology builders, one per model, returning a `jaxley.Cell` and a geometry dataclass describing the per-compartment static parameters;
- `jaxon.stim` — the intracellular and point-source extracellular stimulus helpers, the coupled backward-Euler solver (`extracellular_coupled.py`), the vmapped multi-fiber forward pass (`batch_solve.py`), and the multi-contact ring-cuff field;
- `jaxon.optim` — the recruitment/activation proxy, the selectivity index, and the Adam optimization loops.

The coupled solver integrates the double-cable equation by backward Euler; each time step is a 2 × 2 block-tridiagonal system in the interleaved intracellular and periaxonal unknowns (Fig. 2), solved directly by a JAX-compatible block-Thomas sweep (verified to 5.6 × 10^−11^ mV against a dense LU reference), with back-substitution via `jax.lax.associative_scan` for *O*(log *n*) depth.

**Figure 2.**
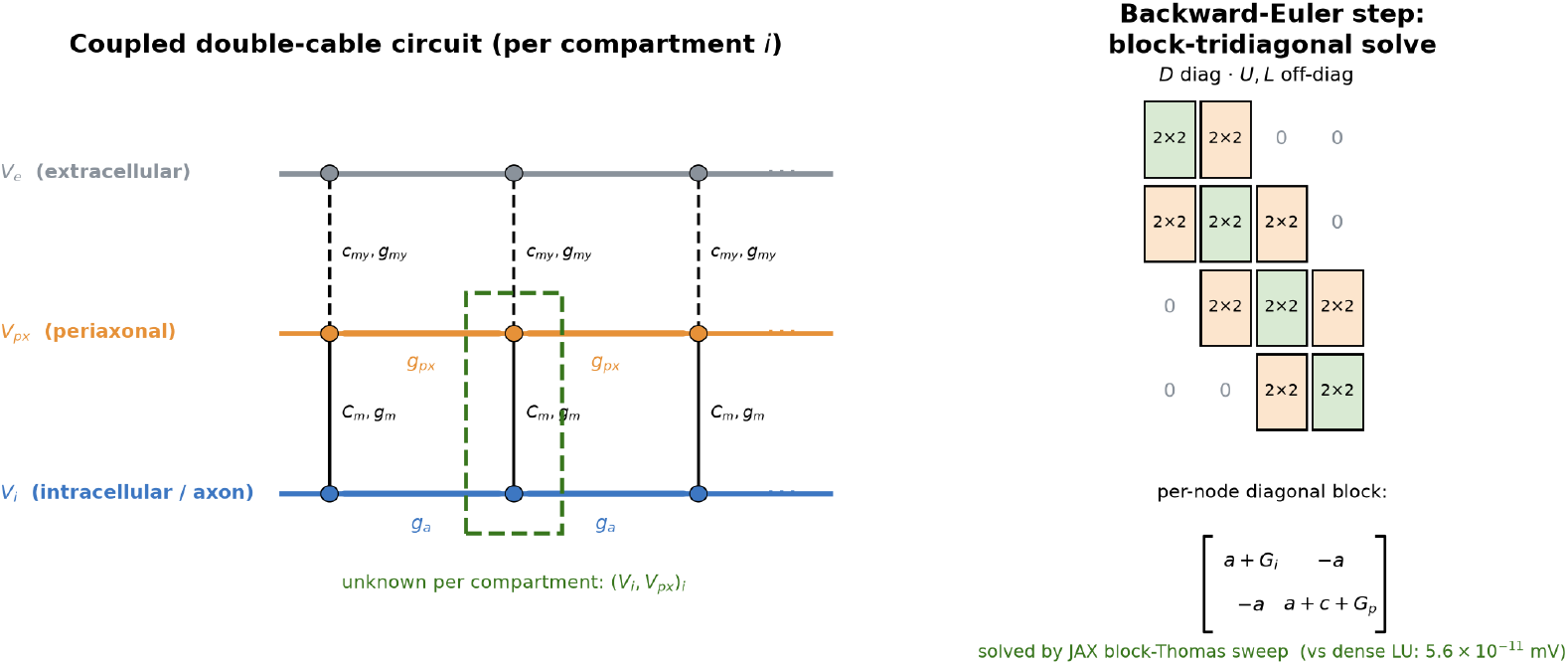
The coupled solver. Left: the double-cable circuit *jaxon* integrates—each compartment couples an intracellular node (*V*_*i*_), a periaxonal space (*V*_*px*_), and the externally imposed extracellular potential (*V*_*e*_, supplied by *golgi* FEM lead fields), through the axolemmal (*C*_*m*_, *g*_*m*_) and myelin (*c*_*my*_, *g*_*my*_) admittances and the axial conductances (*g*_*a*_, *g*_*px*_). Right: each backward-Euler step is a block-tridiagonal system in the interleaved (*V*_*i*_, *V*_*px*_) unknowns, whose 2 × 2 diagonal blocks are solved directly by a JAX-compatible block-Thomas sweep (verified to 5.6 × 10^−11^ mV against a dense LU reference).

Because every operation is a JAX primitive, the same forward pass is (i) batched over fibers with `jax.vmap` for population-scale throughput and (ii) differentiated with reverse-mode autodiff (or a packed finite-difference gradient) for stimulus optimization. For the cohort application, per-contact lead fields are supplied by the *golgi* platform [12, 13] from histology-segmented nerve geometries; a lightweight analytic point-source field is included for local demonstrations.

#### 2.2. Software functionalities

*jaxon*’s major functionalities are:

1. **Canonical fiber models**. Four published axon models—myelinated MRG [3] (with both the original and interpolated parameter tables) and Sweeney [4], and unmyelinated Sundt [5] and Rattay [6] C-fibers— reimplemented as pure-JAX channels and morphologies (Fig. 3).
2. **Coupled extracellular solver**. A backward-Euler integrator for the two-state (*V*_*i*_, *V*_*px*_) double-cable equation—the principled equivalent of NEURON’s `extracellular` mechanism rather than the single-cable activating-function approximation—solved by a JAX-compatible 2 × 2 block-Thomas sweep.
3. **Population-scale batched forward pass**. `jax.vmap` packs *N* fibers into a single vectorized simulation, enabling whole-nerve populations to be simulated and re-scored on one GPU.
4. **Differentiable stimulus optimization**. Three optimization modes: per-contact rectangular amplitudes via a packed finite-difference gradient (*K*+1 configurations in one forward pass); arbitrary *K × T* waveforms via reverse-mode autodiff through the ODE scan (with gradient checkpointing); and joint amplitude-plus-electrode-position search, with the extracellular field recomputed differentiably each step.
5. **Selectivity scoring**. A recruitment (activation) proxy and a selectivity index defined as the difference between the recruited fraction of on-target and off-target fibers, usable both as an optimization objective and as a post-hoc population score.
6. **FEM-field and multi-contact support**. Per-contact lead fields from the *golgi* platform over histology-segmented nerves and a multi-contact ring cuff, with a superposition-based current-steering model; an analytic point-source field is provided for demonstrations and tests.
7. **Validation and benchmarking**. Scripts that reproduce the threshold, conduction-velocity, and propagation-phenomena comparisons against PyFibers-wrapped NEURON, and the CPU/GPU scaling benchmark.

**Figure 3.**
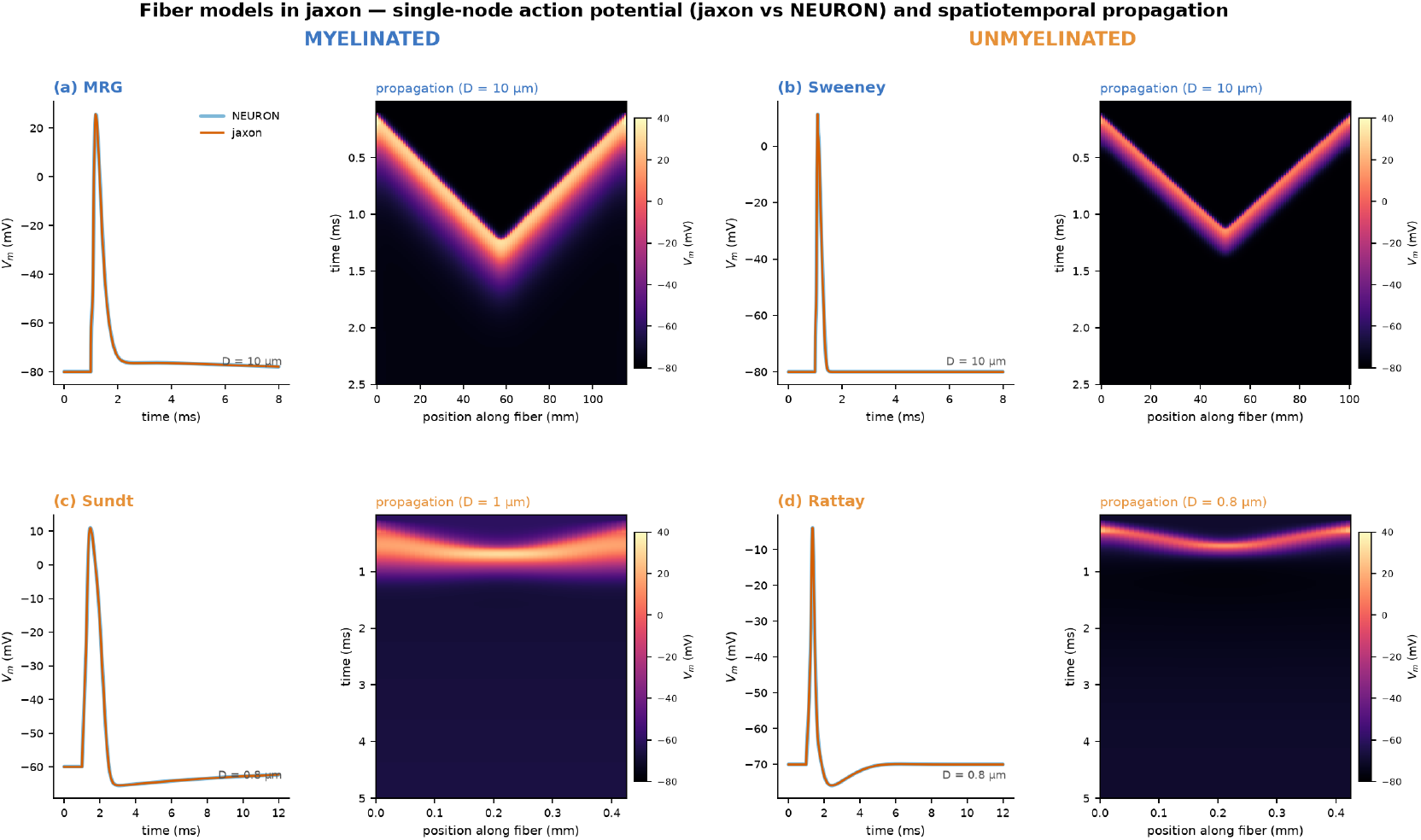
The four fiber models integrated in *jaxon*, grouped by class. For each model, left: the single-node membrane action potential, with *jaxon* (orange) overlaid on the PyFibers-wrapped NEURON reference (blue); right: the spatiotemporal *V*_*m*_(position, time) propagation. The myelinated MRG and Sweeney axons (a, b) show fast saltatory propagation— here two counter-propagating spikes meeting and annihilating in a characteristic collision— while the unmyelinated Sundt and Rattay C-fibers (c, d) propagate slowly and continuously. All curves are *jaxon* output; the NEURON overlay is the validation reference.

#### 2.3. Sample code

The library exposes the model-building, forward-solve, loss, and optimization functions directly, so a study is assembled from a few calls: build one or more fibers with jaxon.fibers (returning a Jaxley `Cell` and a geometry dataclass); assemble the per-fiber static parameters and initial states with `stack_fiber_statics` / `initial_states_batch`; run the batched coupled solve with `batch_integrate_m_max`; and score or optimize with `jaxon.optim` (`selectivity_index`, `run_rect_optimization`). Section 3 gives three worked, runnable examples following this progression.

### 3. Illustrative examples

The three examples below show the typical usage progression—validating one fiber against NEURON, simulating a population in a single batched pass, and optimizing a multi-contact stimulus by differentiating through that pass. All are runnable from the released package; the corresponding end-to-end scripts live in `experiments_v2/`.

*Example 1 — build a fiber and compute an activation threshold*. A single model fiber is built, placed in an extracellular field, and its activation threshold is obtained by the same bisection used for the NEURON comparison. This is the unit that the validation suite runs across 943 configurations.

~~~
import numpy as np
from jaxon.fibers import mrg
from jaxon.stim.batch_solve import (build_fiber_statics,
                                              initial_states_batch,
                                              batch_integrate_m_max)
from jaxon.stim.extracellular import (point_source_potentials_mV,
                                              rectangular_waveform)
dt = 0.005                                    # ms
# Build an MRG fiber (Jaxley Cell + geometry metadata).
cell, geom = mrg.build_mrg(diameter=10.0, temperature=37.0)
# Static point-source profile (unit current) 1 mm above the fiber,
# times a rectangular pulse -> the [T, n_comp] extracellular sequence.
t_grid = np.arange(0.0, 2.0, dt)
profile = point_source_potentials_mV(mrg.section_centers_um(geom),
                                                   src_y_um=1000.0, i0_mA=-1.0) # mV per mA
wave = rectangular_waveform(t_grid, delay_ms=0.1, pw_ms=0.1, amp=0.5) # mA
Ve_seq = wave[:, None] * profile[None, :]    # [T, n_comp]
# Integrate the coupled (V_i, V_px) cable equation; batch_integrate_m_max
# returns the peak Na m-gate, the proxy jaxon uses to decide firing.
fs = build_fiber_statics(geom, dt=dt)
s0 = initial_states_batch([geom])
# … scale ‘amp’ by bisection until the fiber just fires -> threshold.
~~~

*Example 2 — simulate a whole population in one batched pass*. The same forward model, stacked over *N* fibers with jax.vmap, simulates an entire population in a single call. This is the operation that runs at a geometric-mean ∼820× speedup over serial NEURON at *N* = 100,000 fibers, and that makes population-scale re-scoring cheap.

~~~
from jaxon.stim.batch_solve import (stack_fiber_statics,
                                         initial_states_batch,
                                         batch_integrate_m_max)
from jaxon.optim.losses import activation_proxy_batch, selectivity_index
# A population of fibers seeded across the nerve cross-section.
geoms = [mrg.build_mrg(diameter=5.7)[1] for _ in range(n_fibers)]
node_idx = [mrg.node_indices(g) for g in geoms]
fs_batch = stack_fiber_statics(geoms, dt=dt)   # [n_fibers, n_comp, …]
s0_batch = initial_states_batch(geoms)
# Ve_seq_batch: [n_fibers, T, n_comp] -- per-fiber extracellular sequence,
# here from the golgi FEM per-contact lead fields under a chosen stimulus.
m_max = batch_integrate_m_max(fs_batch, s0_batch, Ve_seq_batch, dt=dt)
# Recruitment per fiber, then the population selectivity index.
acts = activation_proxy_batch(m_max, node_idx)
si = selectivity_index(acts, target_mask) # SI in [-1, 1]
~~~

*Example 3 — optimize a multi-contact stimulus by gradient descent*. Because the batched forward pass is differentiable, the per-contact amplitudes of a cuff are optimized directly against the selectivity objective. `run_rect_optimization` drives an Adam loop through the exact cable solve, using the packed finite-difference gradient (*K*+1 configurations per forward pass); an autodiff variant (`run_rect_optimization_autodiff`) differentiates through the ODE scan for arbitrary waveforms.

~~~
from jaxon.optim.optimizer import run_rect_optimization
# Ve_unit: [K, n_fibers, n_comp] -- unit lead field of each of the K contacts.
# pulse_mask: [T] -- the rectangular time course shared by all contacts.
amps, history = run_rect_optimization(
    fs_batch, s0_batch, Ve_unit, pulse_mask,
    node_indices=node_idx, target_mask=target_mask,
    dt=dt, n_steps=90, lr=8e-2,
    amp_clip=(-2.5, 2.5),    # per-contact amplitude bounds (mA)
)
# Re-score the optimized stimulus on the FULL population (cheap in jaxon):
Ve_opt = jnp.einsum(“k,knc->nc”, amps, Ve_unit)[:, None, :] * pulse_mask[None, :, None]
m_max = batch_integrate_m_max(fs_batch, s0_batch, Ve_opt, dt=dt)
print(“population SI:”, selectivity_index(activation_proxy_batch(m_max, node_idx),
                                                             target_mask))
~~~

All three examples are reproducible from the released code; `REPRODUCE.md` documents the full path from a fiber build to every figure and reported number in the companion study [1].

### 4. Impact

*jaxon* changes what is computationally feasible in peripheral-nerve-stimulation modeling along two axes that the reference NEURON stack leaves closed.

First, it **makes population-scale fiber modeling routine**. By expressing the canonical axon models as a `vmap`-batched JAX forward pass, *jaxon* simulates whole fiber populations on a single GPU at a geometric-mean ∼820× speedup over serial NEURON, turning a previously prohibitive whole-nerve re-scoring into a step cheap enough to run inside an optimization loop. This directly enables the central question of the companion study [1]—whether reducing each fascicle to a representative fiber biases predicted selectivity— which was untestable precisely because the population simulation was too expensive.

Second, it **makes the stimulus itself an optimization variable**. Because the whole model is differentiable, per-contact amplitudes, arbitrary waveforms, and electrode positions can be optimized by gradient descent through the exact cable equation, rather than by grid search or a learned surrogate. Unlike a surrogate model, the gradient is of the biophysics, not of a fit to it, so an optimized stimulus can be validated in the same tool that produced it.

Third, it **drops into existing pipelines with minimal friction**. *jaxon* is validated against the community-standard PyFibers/NEURON reference (99.6 % of 943 configurations within 1 %; conduction velocity to machine precision) and released open-source under a permissive license, so it can serve as a gradient-enabled, population-scale replacement for the NEURON forward solver wherever fiber-level thresholds are needed. Its per-contact-lead-field interface connects it to FEM tooling such as *golgi*, positioning it as an inter-operable component rather than a standalone silo.

By combining exact, validated biophysics with GPU-scale throughput and end-to-end differentiability, *jaxon* lets neural-interface researchers ask and answer optimization and population-scale questions that the established serial, non-differentiable tools cannot.

### 5. Conclusions

*jaxon* is an open-source, differentiable, GPU-native reimplementation of the canonical peripheral-nerve fiber models—MRG and Sweeney (myelinated) and Sundt and Rattay (unmyelinated)—built on JAX/Jaxley with a custom coupled (*V*_*i*_, *V*_*px*_) backward-Euler solver. It matches PyFibers-wrapped NEURON on 99.6 % of 943 threshold configurations and to machine precision in conduction velocity, scales to a geometric-mean ∼820× speedup at *N* = 100,000 fibers on one GPU, and exposes exact gradients of the fiber response for stimulus optimization. Validated against NEURON and applied to a species-dependent selectivity question in the companion paper [1], *jaxon* provides a gradient-enabled, population-scale replacement for the NEURON forward solver in peripheral-nerve modeling pipelines.

## Data availability and licensing

*jaxon* is released as open-source software under the MIT license (code meta-data C3/S3). The source code is available on GitHub (https://github.com/CellularSyntax/jaxon), and the tagged v1.0.0 release is archived on Zenodo [14]. The processed data required to reproduce every figure and reported number—the *golgi* -derived per-contact lead fields and the validation, scaling, and cohort-sweep outputs—is archived as a companion Zenodo data record [15] under the Creative Commons Attribution 4.0 International (CC-BY-4.0) license, distinct from the software license. The raw finite-element meshes are *golgi* outputs and are available through the *golgi* platform [12, 13] and the underlying SPARC datasets. Reproduction is documented in `REPRODUCE.md` in the repository.

## CRediT authorship contribution statement

**David Lung:** Software, Investigation, Validation, Writing – review & editing. **Max Haberbusch:** Conceptualization, Methodology, Software, Supervision, Funding acquisition, Writing – original draft, Writing – review & editing.

## Acknowledgments

The authors thank the Medical University of Vienna for access to high-performance computing resources. *(Funding acknowledgments to be completed by the authors.)*

## Current executable software version

**Table 2.**
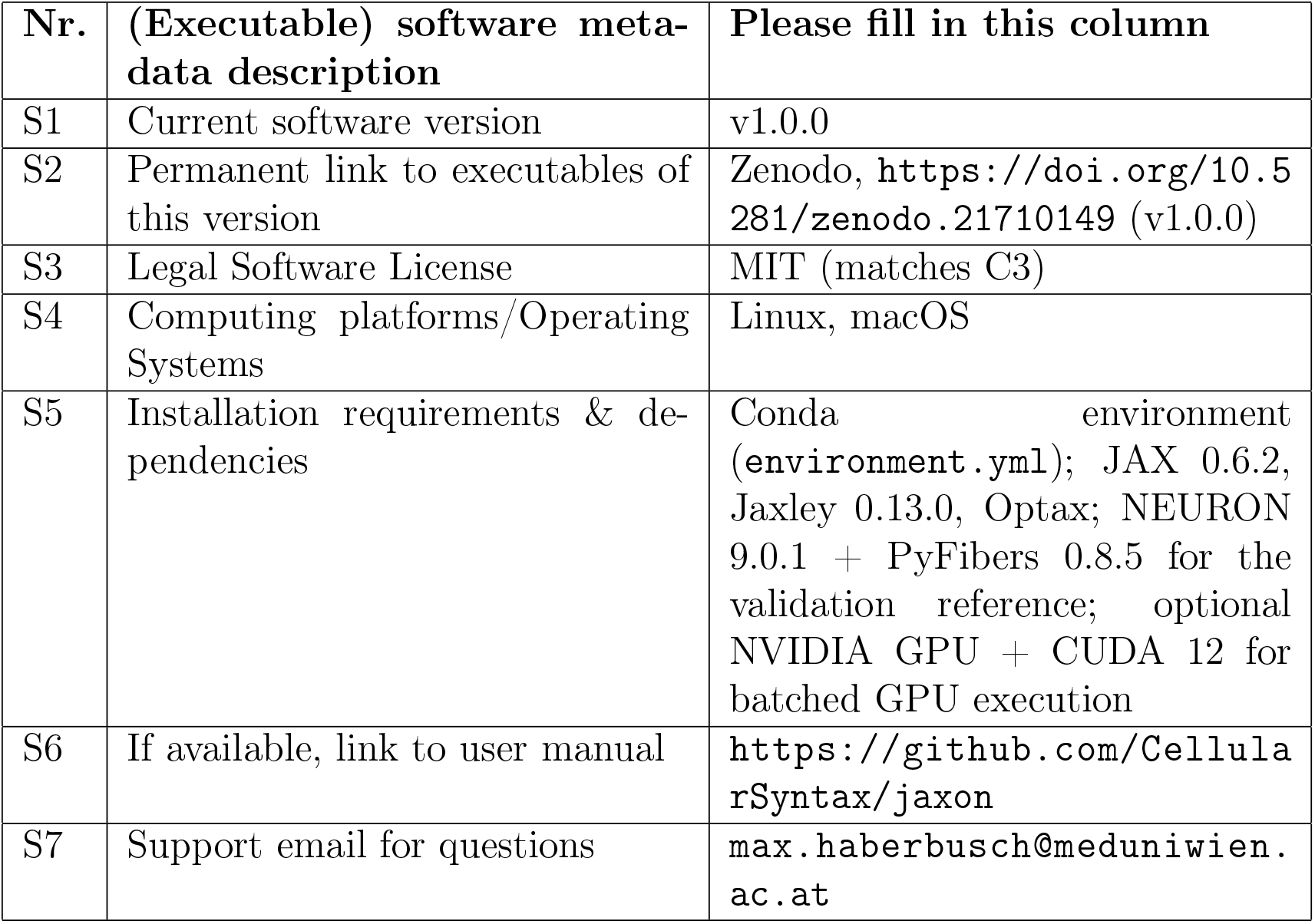
Software metadata.

